# Genetic composition of queen conch (*Lobatus gigas*) population on Pedro Bank, Jamaica and its use in fisheries management

**DOI:** 10.1101/2021.01.07.425691

**Authors:** Azra Blythe-Mallett, Karl A. Aiken, Iris Segura-Garcia, Nathan K. Truelove, Mona K. Webber, Marcia E. Roye, Stephen J. Box

## Abstract

The queen conch fishery in Jamaica is sustained by Pedro Bank, which is the main harvesting site located approximately 80 km south-west from Kingston. Due to its relative size, Pedro Bank has been subdivided into zones for management purposes by the Fisheries Division and the Veterinary Services Division. Understanding whether these sub-divisions reflect different sub-populations is critical for managing exploitation levels because fisheries management must demonstrate that harvesting does not endanger the future viability of the population as queen conch are on Appendix II of the Convention in Trade in Endangered Species of Wild Fauna and Flora (CITES). This determination is essential for the continued export to international markets such as the European Union. Two hundred and eight samples were collected across the entire Pedro Bank and were genetically characterized using nine polymorphic microsatellite loci. Population structure analysis for *Lobatus gigas* from Pedro Bank yielded low but significant values (F_ST_ = 0.009: p = 0.006) and suggested a high magnitude of gene flow indicative of a fit and viable population throughout the bank. Analysis of molecular variance (AMOVA) indicated a 100% variation within individual samples with little variation (0.9%) between populations. In contrast pairwise genetic comparisons identified significant differences between populations located to the south eastern and eastern region of the bank to those in the central and western locations. Bayesian clustering analysis also indicated the likelihood of two population sub-divisions (K=2) on Pedro Bank. The results provided evidence of a weak but significant population structure which has crucial implications for the fishing industry as it suggests the use of ecosystem based management (EBM) in setting quotas to promote sustainable harvesting of *L. gigas* within each monitoring zone on Pedro Bank.

## 1. INTRODUCTION

The queen conch *Lobatus gigas*, (Linnaeus 1758), is an edible marine gastropod mollusk of the order Mesogastropod, and one of six species of the family Strombidae found throughout the Caribbean region [1–5]. With the advent of overfishing in coastal waters along with habitat loss due to coastal and marine ecosystem degradation [6], fishers are travelling further away from traditional fishing grounds to sustain their livelihood and fulfill their need for food security [7,8]. The main conch grounds in Jamaica are located offshore approximately 80 km south-west of the mainland on Pedro Bank. In 1992, Jamaica became the largest queen conch producer in the Caribbean, and the resources on Pedro Bank began to show signs of overexploitation following a pattern of conch population declines elsewhere in the region [9,10].

In response to these events and concerns, the Convention on the International Trade in Endangered Species of Wild Fauna and Flora (CITES), moved queen conch (*L. gigas*) to a listing in Appendix II. This requires that all international trade by signatory countries be accompanied by a valid CITES permit, issued by the authorized National CITES Management Authority and endorsed by the Scientific Authority [2,11,12]. Mahon et al. [9], predicted a collapse in the Jamaican Conch Fishery within three years if adequate management measures were not put in place to reduce or eliminate the harvesting of immature individuals. This led to the drafting of the Conch Management Plan and between 2008 and 2011, conch production decreased from the unsustainable early 1990s harvest figures of close to 3,000 mt to around 400 mt, with a value of approximately 2.6 million US dollars [7,10]. This decrease can be attributed to the implementation of management strategies by the Ministry of Industry, Commerce, Agriculture and Fisheries (MICAF) to control exploitation of dwindling stocks. Some of the management regulations currently in place include, licensing of conch fishers and conch fishing vessels, closed season, transferable quota system based on National Total Allowable Catch (NTAC), minimum weight of processed meat, vessel monitoring system (VMS), fishing effort report on each fishing trip along with inspection of catch on landing and a conch export levy which is put into a fisheries management fund for the development of a sustainable fisheries sector in Jamaica [2,13–15].

Pedro Bank (Fig 1A), has an area of 8040 km^2^, a maximum length of 168 km and a maximum width of 83 km in the west [16]. It consists of extensive sand flats interspersed with patches of macroalgae made up of various species, rubble, scattered patch reefs and shoals [10,17,18]. Due to its size, fisheries management zones were assigned across the bank, based on depth and fishing pressure (Fig 1B), by Jamaica’s Veterinary Services Division (VSD) which is the competent authority in charge of monitoring conch production areas. These demarcations have allowed for more effective monitoring of the main conch production area as fishing activities can be prohibited based on the health of marine gastropods and prevailing sanitary conditions within a specified zone [19]. Genetic diversity assessments and comparisons between harvesting zones may help assure managers of the integrity of the fishery products if a harvesting zone is closed and allow for specific management measures to be applied to each zone especially where sub-populations are found. The identification of sub-populations could then effect fishing effort control to a specific area and bring management to the sub-population level. In addition, the information on population genetic structure can also be used in the design of stock enhancement programmes by ensuring appropriate genetic diversity in each area [20]. This is especially useful as selection may favor different genotypes from different habitats and geographic locales [21].

**Figure 1.**
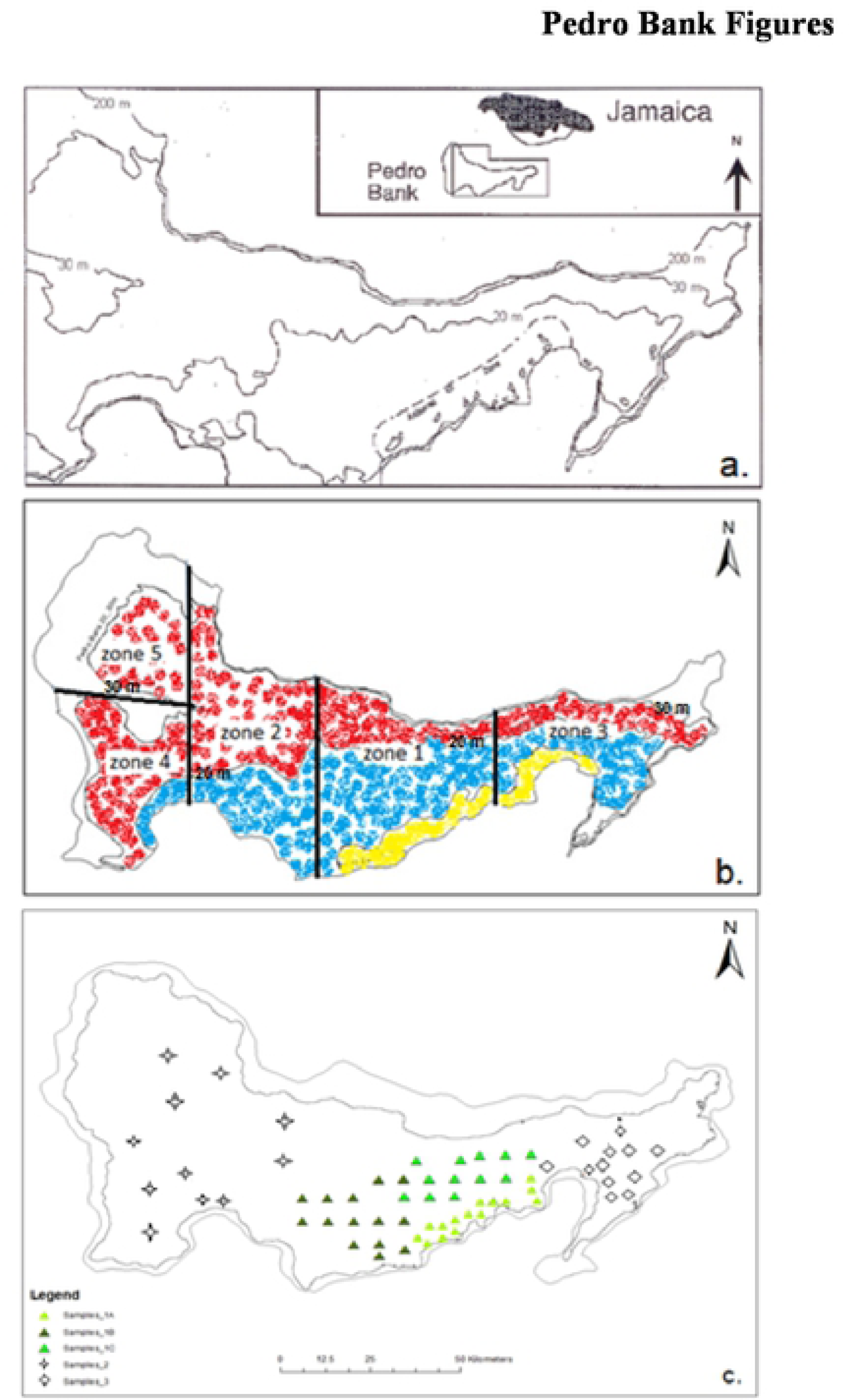
Map of Pedro Bank: (a.) Highlighting location approximately 80 km SW of mainland Jamaica with major contour lines demarcating Artisanal zone (0-10 m) and Industrial Fishing zone (11-20 m) (Adopted from [11]). (b.) VSD Monitoring zones 1-5 established across the bank for management purposes and straddling the depth zones 0-10 m in yellow (artisanal zone), 11-20 m in blue (industrial zone) and the 21-30 m zone in red. (c.) Sampling locations within monitoring zones. Three sets of samples were located in zone 1 as differentiated by shades of green triangles. Diamond shapes were representative of samples in zone 3 while zones 2, 4 and 5 were grouped into one sample set (star shapes). Map produced using ArcGIS ver. 10.3.1

Providing more effective monitoring of the Pedro Bank through genetic labelling of conch will also strengthen international trade by meeting stringent requirements set out by the European Commission Decision [19] which outlined special conditions for the import of marine gastropods originating in Jamaica, the U.S. Food and Drug Administration (FDA), Seafood Guidance Documents and Regulatory Information for Shellfish Programs and the *Codex Alimentarius* code of practice for fish and fishery products. The aim of this study was to detect fine-scale population structure and how that could impact long term conservation and sustainable harvest of *L.gigas* stocks on the Pedro Bank along with traceability of product from harvesting site to market.

## 2. METHODS

### 2.1 Sample collection and processing

A total of 208 *L. gigas* individuals were sampled across the five monitoring zones on Pedro Bank (Fig 1C), with 144 samples collected from the 2015 conch abundance survey (Fig 2) and an additional 64 samples through monitoring by the competent authority of MICAF, (Table 1). A tissue sample of approximately 4 cm^2^ was removed from the tip of the mantle, rinsed in clean water to remove excess mucus and dirt, then placed on blotting paper for drying before extraction (Fig 3). DNA for each individual sample was extracted and purified using the AutoGenprep 965, a fully automated high throughput system utilizing the solution phase organic extraction method at the Smithsonian Institution’s Laboratory for Analytical Biology National Museum of Natural History (LAB-NMNH). The samples were returned as pellets to be re-suspended with 50 ml Elution Buffer before the DNA quality and quantity was assessed using gel electrophoresis and NanoDrop 1000 Spectrophotometer (Thermo Fisher Scientific).

**Table 1.** Samples collected from the five monitoring zones across Pedro Bank.

| Sample Locations | Monitoring Zones | Number of Samples |
| --- | --- | --- |
| 1A | 1 | 46 |
| 1B | 1 | 40 |
| 1C | 1 | 42 |
| 2 | 2,4,5 | 34 |
| 3 | 3 | 46 |
| **Total** |  | **208** |

**Figure 2.**
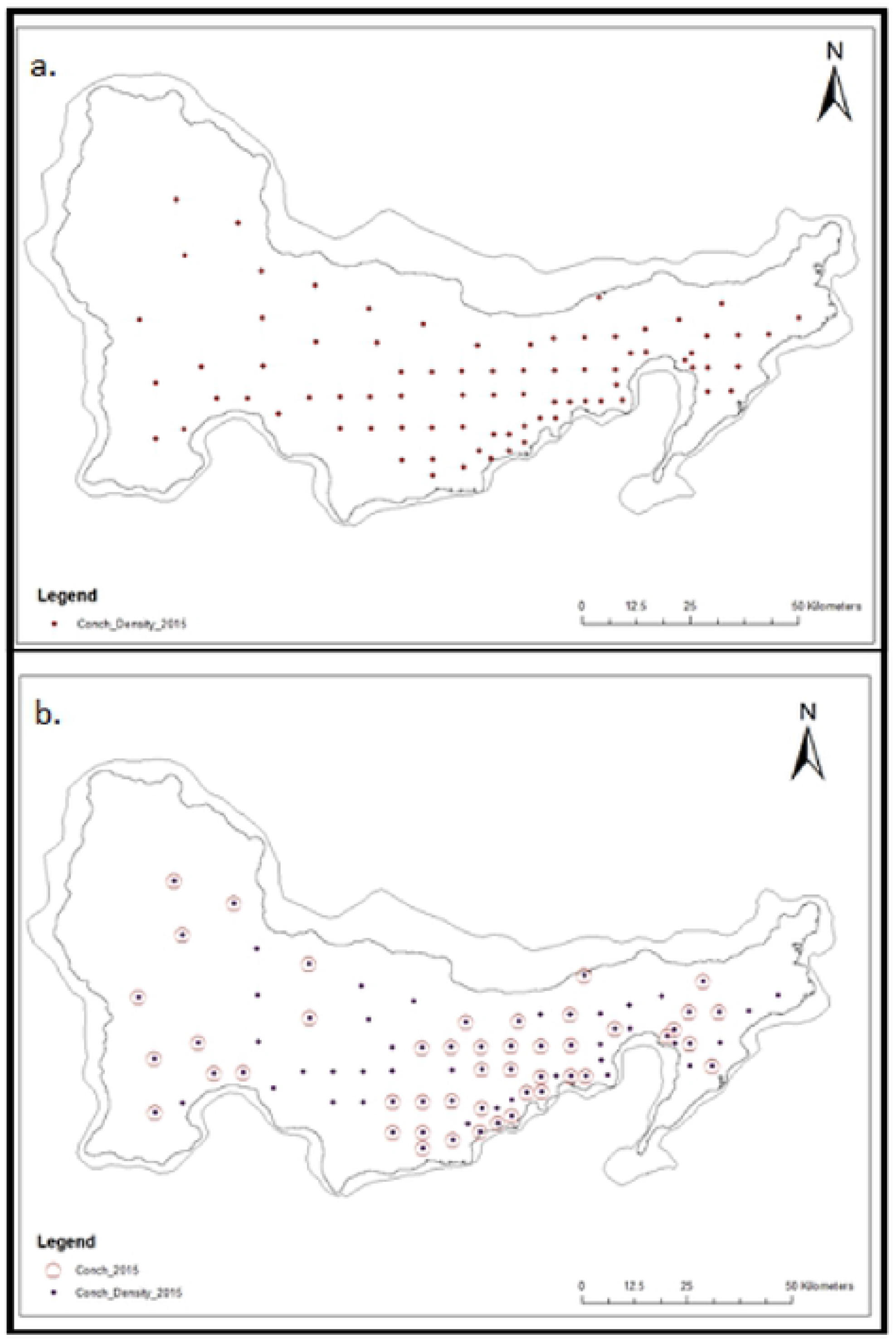
Sampling locations on Pedro Bank. (a.) Sites sampled in the 2015 conch abundance survey. (b.) All 80 sites from the 2015 survey with 47 highlighted to show samples taken for genetic studies in keeping with the 1:2:1 ratio associated with fishing pressure across all 3 depth zones while allowing for maximum coverage of Pedro Bank. Map produced using ArcGIS ver. 10.3.1

**Figure 3.**
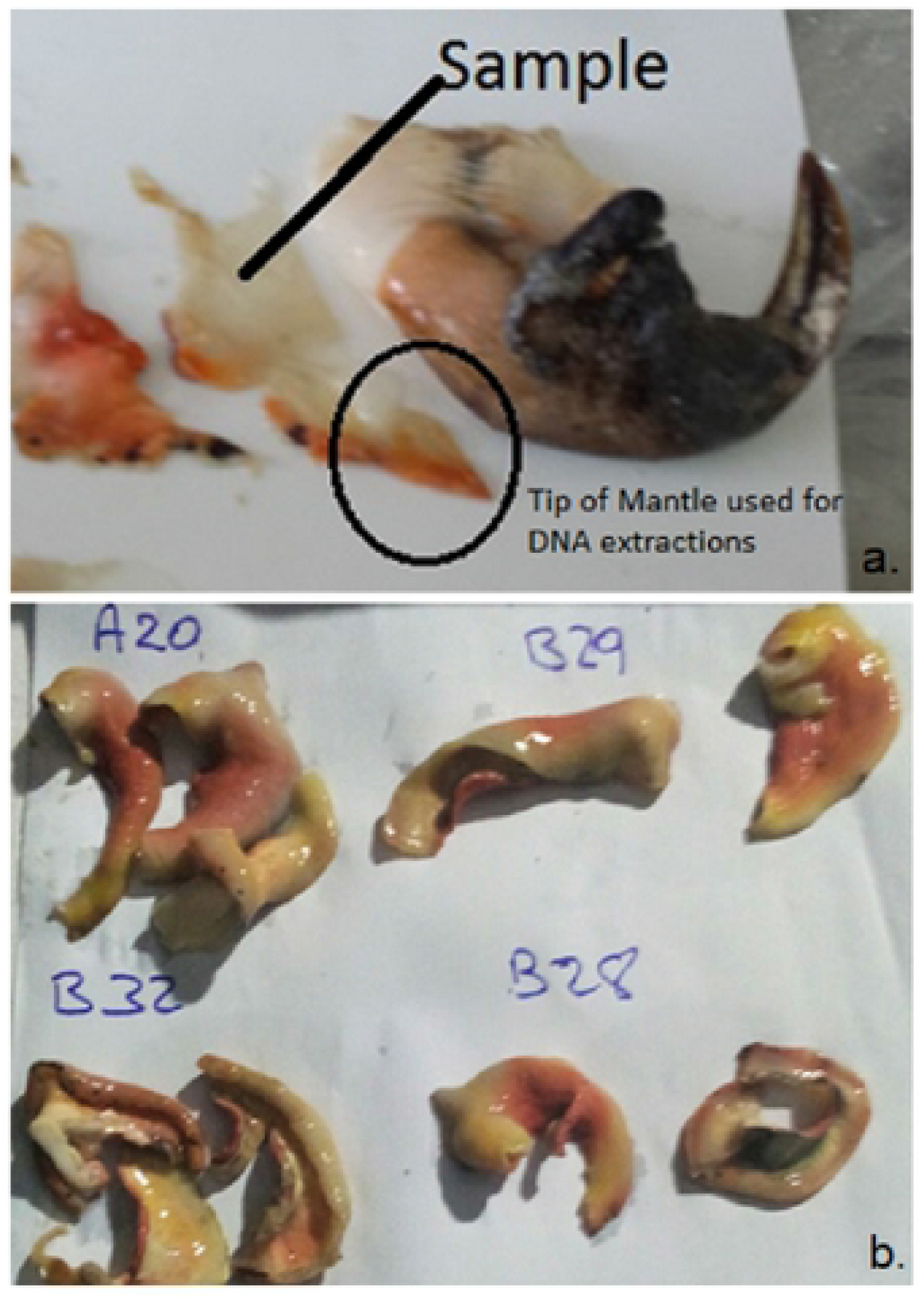
Individual conch samples: (a.) Showing tip of mantle collected for genetic analysis. (b.) Air dried on paper with coded sampling location from zones on Pedro Bank prominently displayed. (Photographs by author)

### 2.2 Microsatellite genotyping

All conch individuals were genotyped for nine microsatellite loci: Sgig1, Sgig2 and Sgig6 [22], and ConchPR11F, ConchF29, ConchF17, ConchPR1F, ConchF21 and ConchF23 [23]. These loci were successfully grouped into 3 multiplexes (Table 2), and amplified using the Qiagen Type-it^®^ Microsatellite PCR kit in a 10μL reaction following the standard protocol. The amplification was performed on a Dyad Thermocycler (Bio-Rad) under the following PCR conditions: 95°C for 5 minutes denaturation, followed by 28 cycles of 95°C for 30 seconds, annealing at 62°C for 90 seconds for MP2 and 63°C for MP1 and MP3, 72°C for 30 seconds and a final extension step at 60°C for 30 minutes. PCR products were separated on an ABI 3730xl DNA analyzer (Applied Biosystems) with ROX size standard and scored manually using GeneMapper^®^ Software v 3.7 (Applied Biosystems). All genotyping was conducted at LAB-NMNH.

**Table 2.**
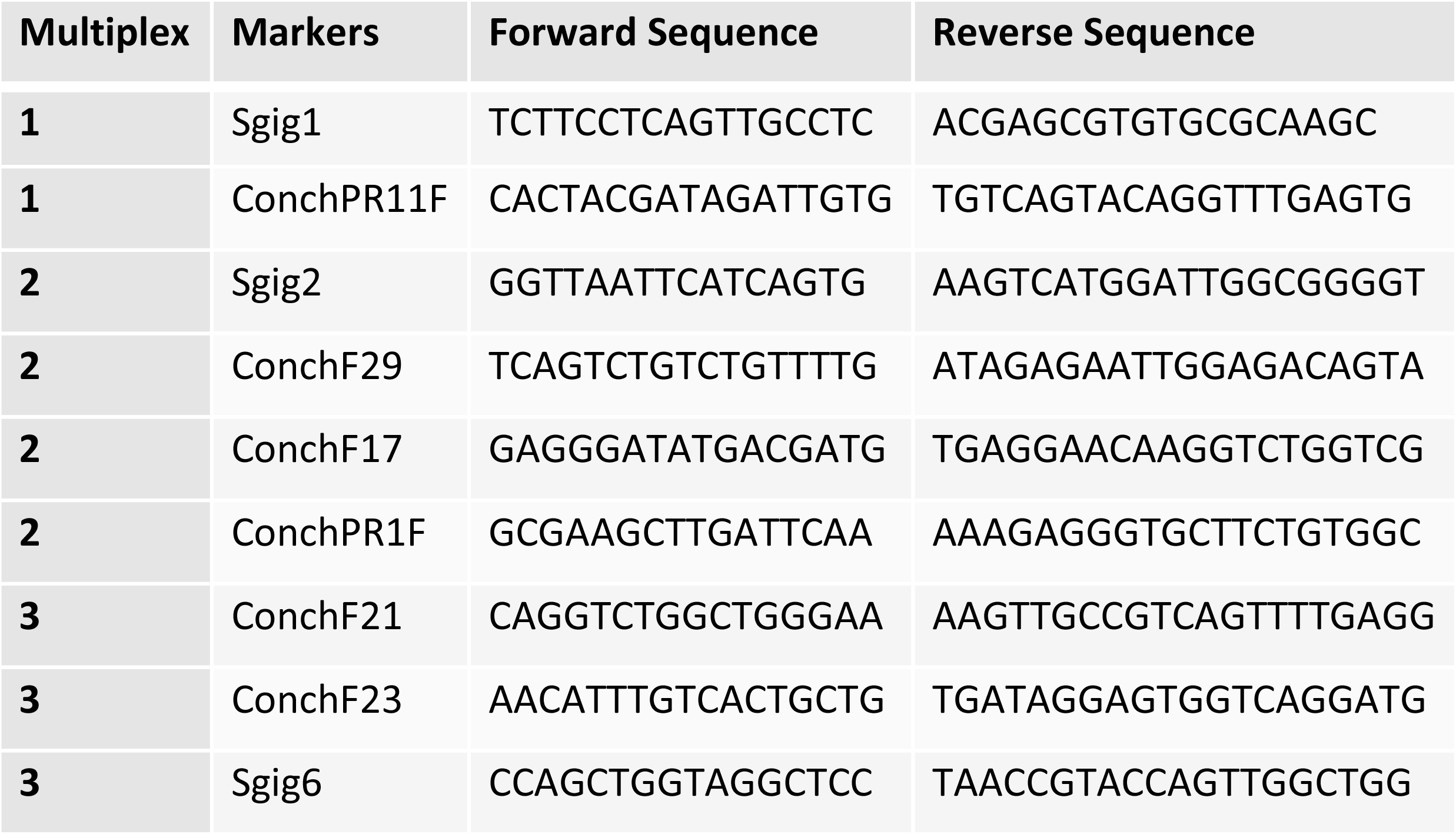
Multiplex grouping for microsatellite markers and primer sequence.

Irregularities in the data set such as genotyping errors caused by the presence of null-alleles and scoring errors due to large allele drop-outs and stuttering were tested for using Micro-Checker version 2.2.3 [24]. FreeNA was used to estimate genetic distance for each population pair with and without the ENA (excluding null alleles) correction [25]. Linkage disequilibrium among all loci and deviations from Hardy-Weinberg Equilibrium (HWE) were calculated at each locus for all populations sampled using the genetic software Arlequin version 3.5.2.2 [26] and Genepop version 4.2. [27,28].

Estimates of genetic diversity, including total heterozygosity estimated from observed heterozygosity (Ho) and expected heterozygosity (He), number of alleles per locus, effective number of alleles along with variance in allele frequency (*F_ST_*) based on 1,000 permutations were computed in Arlequin version 3.5.2.2 [26]. Relatedness was explored by calculating the inbreeding coefficient within individuals at each locus (*F_IS_*) and the inbreeding coefficient adjusted for bias (*G_IS_*) using GenAlEx 6.502 with probability data based on 999 permutations [29,30].

The level of genetic differentiation within and between populations was estimated using the analysis of molecular variance (AMOVA), and the hierarchical AMOVA was effected among populations using Arlequin ver 3.5.2.2. Pairwise population matrices were calculated using the G-statistics option in GenAlex 6.502., with probability based on 999 permutations. To determine the most probable number of populations clusters (K) and assign each individual sampled to these clusters based on their genotype, a Bayesian assignment method was performed as implemented in STRUCTURE version 2.3.4 [31]. A total of 1,000,000 MCMC (Markov Chain Monte Carlo) repetitions were performed with a burn-in period of 10,000 iterations for K ranging from 1-10 in consideration of the maximum number of expected populations, with 10 independent runs for each putative number of populations. The most likely number of clusters was inferred for population subdivision by the modal value of ΔK, corresponding to the rate of change in the likelihood of K following recommendations by Evanno et al. [32]. DARwin (Dissimilarity Analysis and Representation for Windows) version 6 software [33] was used to visualize the genetic relationship between individuals within and between populations by using the weighted neighbour joining method [34].

## 3. RESULTS

Linkage disequilibrium was evident in three pairwise comparisons between loci using Fisher’s method (Sgig 1 and ConchF23, Sgig 1 and Sgig 6, ConchF23 and Sgig6). After false discovery rate (FDR) correction (p < 0.001) only comparisons with Sgig 2 appeared significant. The results showed two significant deviations from Hardy Weinberg Equilibrium (HWE) from different populations (1C and 3), and at different microsatellite loci (Sgig2 and ConchF23) respectively. Analysis in Micro-Checker [24] suggested that these deviations were caused by the presence of null alleles as there was no consistent evidence of scoring errors due to stuttering or evidence of large allele dropout across all loci. The analysis with FreeNA using the ENA method to correct for bias induced by the presence of null alleles [25], indicated that calculations of Global *F_ST_* = 0.0084 and a corrected Global *F_ST_* = 0.0083. GenAlEx computations of G-statistics such as *F_ST_* adjusted for bias *G_ST_*, and standardized *G”_ST_* which further corrected for bias when number of population is small (S1 Appendix A), showed values which correlated with calculated *F_ST_*, therefore all microsatellite loci were included in further analysis and reported using the fixation index.

All nine microsatellite loci were highly polymorphic across all five sampled locations on Pedro Bank. The average number of alleles per locus ranged from 12.4 to 15.1 with 6.7 (± 0.79) being the mean effective number of alleles across all sites (Table 3). Zone 1 which constitutes sample locations 1A, 1B and 1C displayed the highest numbers of private alleles and the highest percentages of genetic diversity which was 78% on average across all sampled locations. Average observed homozygosity was slightly higher than expected heterozygosity, 0.81 and 0.77 respectively across Pedro Bank with the inbreeding coefficient F_IS_ detected at low levels and was negative at three of the five locations sampled.

**Table 3.** Genetic diversity across sampling locations on Pedro Bank. Na: average number of alleles per locus, Ne: number of effective alleles per locus, Np: number of private alleles, He: expected heterozygosity, Ho: observed heterozygosity, Ht: total heterozygosity and Fis: inbreeding coefficient

| Sampling Locations | Na | Ne | Np | He | Ho | Ht | Fis |
| --- | --- | --- | --- | --- | --- | --- | --- |
| 1A | 15.111 | 7.536 | 1.444 | 0.779 | 0.801 | 0.783 | 0.003 |
| 1B | 14.556 | 7.210 | 1.556 | 0.799 | 0.850 | 0.804 | -0.046 |
| 1C | 14.222 | 6.891 | 1.222 | 0.792 | 0.794 | 0.796 | 0.040 |
| 2 | 11.444 | 5.462 | 0.667 | 0.777 | 0.846 | 0.782 | -0.067 |
| 3 | 12.444 | 6.562 | 0.778 | 0.736 | 0.768 | 0.736 | -0.014 |
| **Mean** | 13.556 | 6.732 | 1.133 | 0.776 | 0.812 | 0.780 | -0.017 |
| **SD** | 1.546 | 0.797 | 0.396 | 0.024 | 0.035 | 0.026 | 0.042 |

Analysis of molecular variance (AMOVA) indicated a 0.9% genetic differentiation among sampling locations, while variation within individuals was 100%. The hierarchical AMOVA (Table 4) indicated a low but significant population variation (*F_ST_* = 0.009: p = 0.006). Pairwise *F_ST_* comparisons ranged from 0.007 to 0.15, with four significant comparisons observed after FDR correction (Table 5), three of which included sampling location 3. The genetic differentiation between sampling locations was consistent with dendrogram constructed using the DARwin software Neighbour-Joining method (Fig 4) and the principal component analysis (PCA) scatterplot (S2 Appendix B) which showed a high degree of overlap between sampling locations.

**Table 4.**
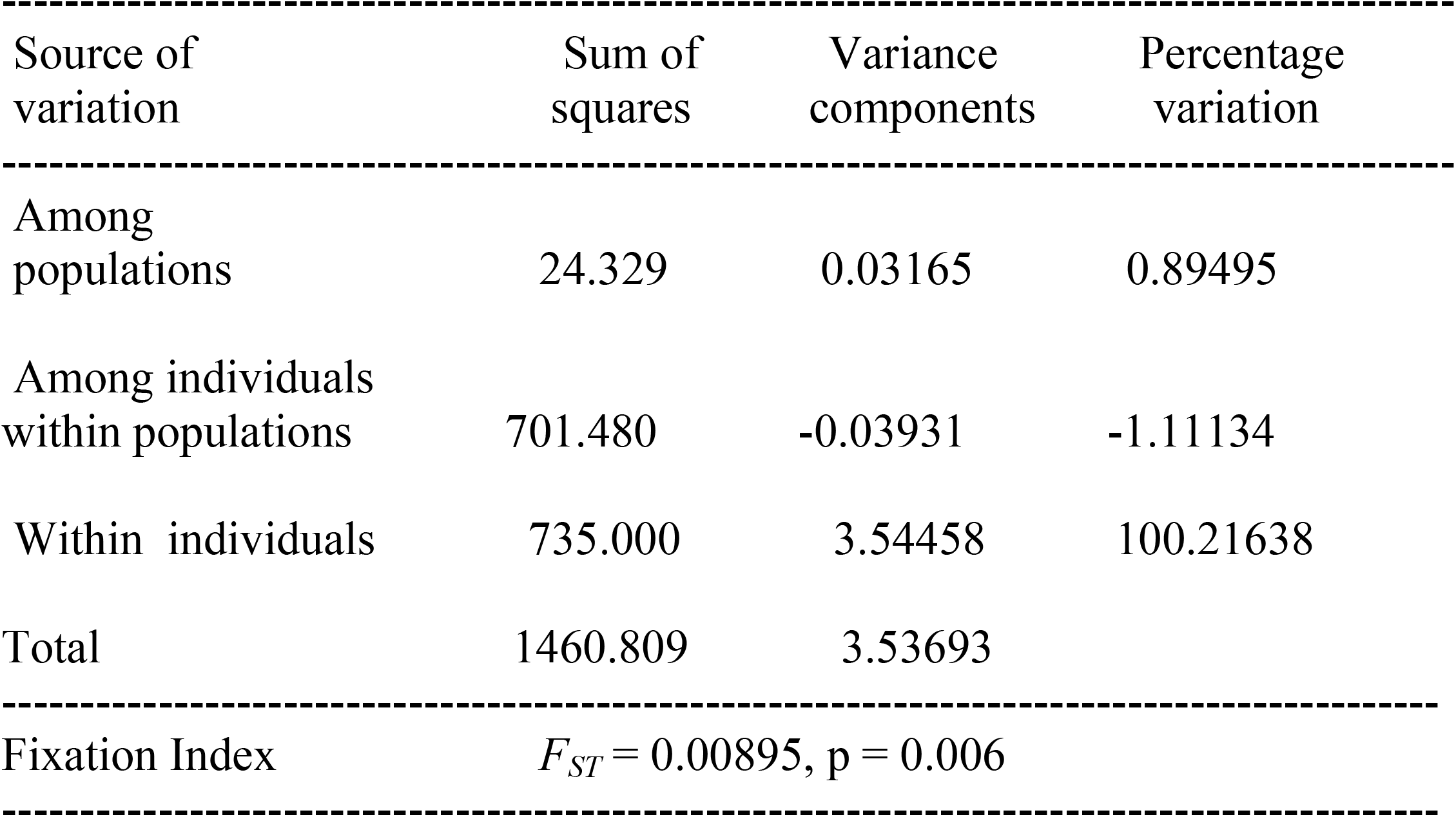
Global AMOVA results as a weighted average over all nine loci across the five locations sampled.

| Source of variation | Sum of squares | Variance components | Percentage variation |
| --- | --- | --- | --- |
| Among populations | 24.329 | 0.03165 | 0.89495 |
| Among individuals within populations | 701.480 | -0.03931 | -1.11134 |
| Within individuals | 735.000 | 3.54458 | 100.21638 |
| Total | 1460.809 | 3.53693 |  |
| Fixation Index | $F_{ST} = 0.00895$, $p = 0.006$ | | |

**Table 5.** Pairwise comparisons of the five populations sampled across Pedro Bank. *F_ST_* values below the diagonal and p values above. Significant values highlighted.

|  | 1A | 1B | 1C | 2 | 3 |
| --- | --- | --- | --- | --- | --- |
| 1A | 0.000 | 0.094 | 0.172 | 0.001 | 0.050 |
| 1B | 0.007 | 0.000 | 0.806 | 0.023 | 0.001 |
| 1C | 0.007 | 0.005 | 0.000 | 0.037 | 0.003 |
| 2 | **0.015** | 0.010 | 0.010 | 0.000 | 0.001 |
| 3 | 0.007 | **0.012** | **0.010** | **0.021** | 0.000 |

**Figure 4.**
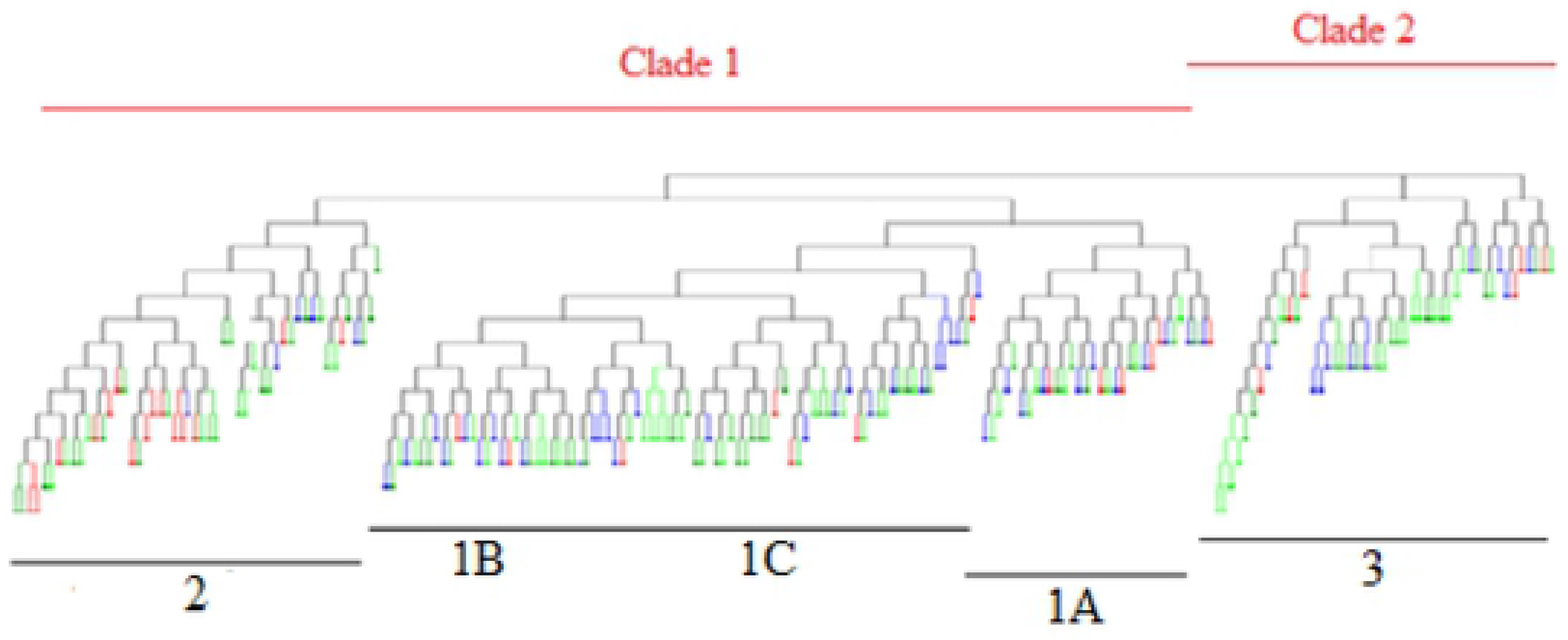
DARwin Neighbour-joining tree showing the admixture of 208 individuals based on estimates of average distances from nine polymorphic microsatellite loci used to characterize the genetic population of *Lobatus gigas* across Pedro Bank. Each location sampled was assigned a colour which magnifies the level of admixture of individuals within and between populations. Below the dendrogram are four defined groups of individuals, while two clades are highlighted above indicating the most likely populations among the samples.

The optimum number of sub-populations (K) of queen conch on Pedro Bank based on the largest value of ΔK using the Bayesian clustering analyses performed in Structure was K=4 (Fig 5). There was extensive admixture of individuals across the Pedro Bank, hence the clusters as shown in the bar plot were not well defined (Fig 6), and showed a high likelihood of K=2, supported by the dendrogram (Fig 4) which also displayed an admixture of individuals from the different locations sampled. The two clusters identified comprises cluster 1 (sampling locations 1A, 1B, 1C and 2), situated to the centre and western end of Pedro Bank and cluster 2 represented by sampling location 3 located on the eastern side of the bank. These results indicating high levels of relatedness was also reflected by the population assignment tests performed in GenAlEx which indicated 68% of individuals being assigned to the population in which it was sampled while 32% indicated a likelihood of being assigned to other populations (S3 Appendix C).

**Figure 5.**
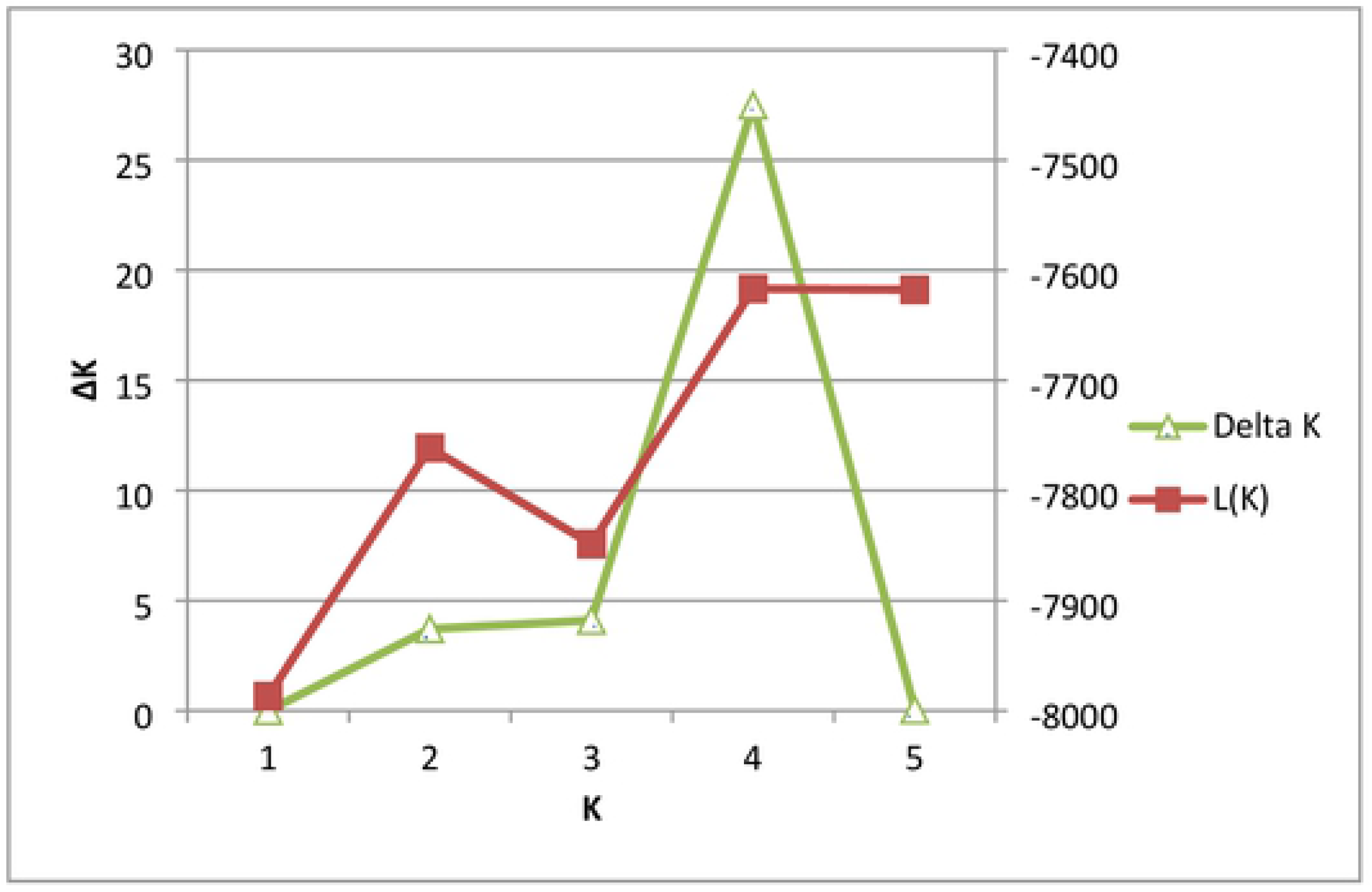
Determination of homogeneous populations (K) using the modal value of an ad hoc quantity (ΔK) which is based on the second order rate of change in the likelihood function [32].

**Figure 6.**
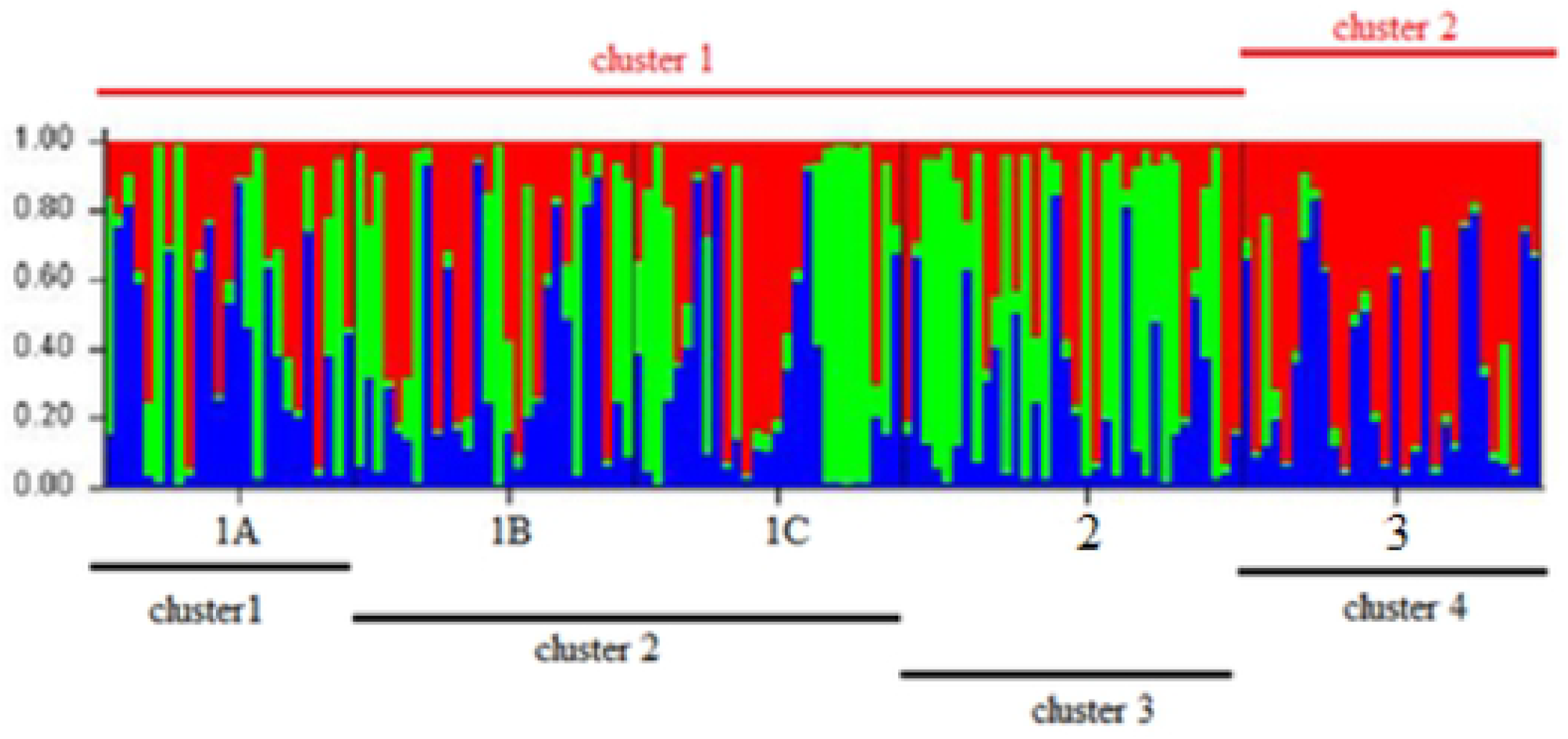
Bayesian clustering analysis of *Lobatus gigas* samples from five sampling sites across Pedro Bank as performed using STRUCTURE. Samples were assigned to four (K=4) population clusters represented by the lines below the bar graph. Lines above the bar plot represents the most reliable populations from the reassignment of individuals as reflected using samples collected from the Queen conch survey only (S4 Appendix D).

## 4 DISCUSSION

This is the first genetic study where all monitoring zones encompassing different depth strata on Pedro Bank were sampled for genetic assessment. Despite the heavy exploitation of queen conch on Pedro Bank, the high genetic diversity values observed within each group of *L. gigas* sampled, is indicative of a vigorous population. This high genetic diversity is likely due to the large area represented by the bank resulting in spatial variability across the Pedro Bank with little population isolation. These results are expected in large populations, which is consistent with the high conch density 374 conch/ha reported by Morris et al. [35] from the Pedro Bank queen conch 2015 abundance survey. Each group exhibited high heterozygosity for all loci with an average of 0.77(± 0.024), signifying high genetic variability across the bank. Despite the occurrence of high genetic variability maintained by gene flow within the Pedro Bank population, Mitton et al. [36] cautioned against making the assumption that conch is a randomly mating population as they identified gaps in allele frequencies from the populations studied within the Caribbean region. Slatkin [37] also observed that gene flow in natural populations varied widely among species and is dependent on the method of analysis used, one such method being the use of private alleles.

Private allelic richness can be used as a simple measure of genetic distinctiveness as it has proven to be informative in conservation genetic studies [38,39]. Slightly elevated frequencies of private alleles were seen in sampling locations 1A, 1B and 1C corresponding to monitoring zone 1 (Fig 1B), which is the most exploited zone. Increased harvesting of species can lead to the depletion of genetic information and later extinction and recolonization of entire populations. Whilst this may prevent differentiation of sub-populations within an area [37,40], an increase in private allele frequency can also be attributed to inbreeding due to the cumulative effect of genetic drift [41]. However, due to the difficulties in collecting data on alleles that are not found in all subpopulations, F-statistics is a better method by which gene flow can be estimated in natural populations [42].

Our results (Table 3) indicated a slightly higher value for average observed homozygosity (Ho) than expected heterozygosity (He). Increased homozygosity is an indication of inbreeding which occurs as a result of mating between two related individuals in a population [43,44,45], or as a result of severe reduction in population size from exploitation [46]. Positive inbreeding coefficient, albeit low values (F_IS_ = 0.003 and 0.040) were observed in the most heavily fished areas suggesting some level of relatedness due to removal of alleles from the population. However, the average inbreeding coefficient across Pedro Bank was negligible signifying a population with high genetic variability, which is less related than is expected under the assumption of random mating. Inbreeding can contribute to the failure of a species to recover from exploitation as it increases the frequency of deleterious alleles [44,47]. Despite this detrimental effect, inbreeding has been used for decades to increase favourable selection responses leading to improvements and ensuring uniformity within livestock breeds [35,43] and to promote economically important traits such as fast growth in food fish production [48]. In fact Gjedrem and Robinson [49] noted that although significant strides have been made within the last 40 years to manage genetic diversity and conserve fish populations, genetic improvements from selective breeding in aquatic animals is far behind that achieved for terrestrial animals and plants.

Kenchington and Heino [50], regarded selective fishing as a possible means of not only affecting life history traits such as age and maturation, but as a means to influence the dynamics of fish populations and ultimately sustainable yield of a population. This observation was also made by Stoner et al. [51] when they reported that the selective removal of larger individuals by fishing, had significant effects on density and biomass as negative growth rates can occur within the population if density was <56 adult conch ha^−1^ [52] or below the recommended 100 conch ha^−1^ [7]. The high conch density calculated for Pedro Bank in the 2015 queen conch abundance survey, especially in the south and south-eastern sections corresponding to monitoring zone 1 of the study area, provided no support for impending local extinction despite heavy exploitation. In fact, the numbers suggest the existence of a substantially large population with high genetic variability [27].

Low estimates of population structure are typically found in benthic marine invertebrates such as *L. gigas* with an extended planktonic larval stage, which may last up to 8 weeks and hence have a high dispersal potential [4,53,54]. Our estimates of population structure indicated a weak but significant population variation (F_ST_ = 0.009: p = 0.006), across Pedro Bank, suggesting genetic homogeneity which can be maintained by the exchange of genetic material between a few individuals per generation [36,37,55]. Significant genetic differences were seen among groups and within individuals but not among individuals within each group sampled. This pattern of genetic variation over small spatial scales was described by Campton et al. [54] as genetic patchiness which can be attributed to post-settlement natural selection where variation among groups is greater than variation within each group. Similar patterns of genetic structure was observed in previous studies of *L. gigas* within the Caribbean region by Mitton et al. [36] and Kitson-Walters et al. [56], and for *Stegastes partitus* (damselfish) by Villegas-Sanchez et al. [57]. The latter attributed the small-scale genetic variation to a temporally unstable genetic structure, a phenomenon known as ‘sweepstake-chance effect’ where most recruits to the fishery are from a few individuals due to random assortment and matching of reproductive activity with oceanographic conditions conducive to spawning, fertilization, larvae development and recruitment.

While many of these processes may contribute to the fine-scale and large-scale spatial variation within the population on Pedro Bank, the results from the pairwise comparisons which indicated that the easternmost group (sampling location 3) was the most significantly different from the other groups in the central and western end, supports the mechanism of genetic variation caused by oceanographic conditions. Surface currents in the Caribbean generally flow towards the west [50,58,59], and Jamaica is supplied from large upstream reef areas and as such may be less susceptible to overfishing and species loss [53,58]. Kitson-Walters et al. [56] also suggested that Pedro Bank receives recruitment from outside the Jamaican exclusive economic zone (EEZ) corroborating the hypothesis that queen conch recruitment would be greatest on the eastern end of the bank, which also has the highest conch density [27]. The substrate cover in that area is predominantly sand sediment, seagrass, coral reefs and small areas of macroalgae [18,60]. Previous studies conducted on Pedro Bank indicate a direct relationship between substrate type, depth and *L. gigas* density [2,11,18, 27,61]. Tewfik and Appeldoorn [61] and Morris [18], found that substrate type and depth were influential in the distribution of queen conch across Pedro Bank, while Stoner et al. [62] reported habitat limitation as the most plausible explanation for low abundances in the conch population west of Cat Island in the Bahamas.

The neighbour joining (NJ) tree is in congruence with the pattern of genetic structure found (Fig 4), where Group 3 was significantly different from the other groups except for Group 1A which is closest geographically and consistent with the genetic structure pattern of queen conch in Yucatan, Mexico [20]. Contrastingly, Campton et al. [54] described queen conch as displaying spatial and temporal genetic variation as geographical distance did not correlate with relatedness of individuals, instead he attributed this genetic patchiness to pre-settlement events due to stochastic variations in the marine environment, temporal variations in recruit source for each locality and/or genetic drift. The distribution of queen conch on Pedro Bank was also described as patchy by Appeldoorn [11] due to densities found throughout the bank from sporadic larval settlement which may also lead to distinct genetic populations being formed [14]. A combination of barriers to dispersal and ocean circulation could also be used to explain variation between groups of marine species [63].

The Bayesian clustering analysis indicated four clusters (K=4) with an admixture of individuals assigned to all clusters indicative of mixed ancestry with a richer and more complex representation than is traditionally assumed by clustering models where each individual belongs to a unique cluster [64]. This analysis also indicated a high likelihood at (K=2), which corresponded to results obtained using samples collected only from the 2015 queen conch abundance survey (S4 Appendix D) which had less individuals to assign. Kalinowski [65] concluded from his simulation exercise that clusters produced by the software STRUCTURE can be strongly influenced by differences in sample size but the same data set can produce realistic results when using traditional methods such as pairwise comparisons. The most probable number of populations identified on Pedro Bank was two which corresponds to the pairwise comparisons where spatial variation was evident between sample location 3 and the rest of the bank.

Sample location 3 situated on the eastern side of Pedro Bank (zone 3) has a predominant depth of 11-20 m and considered to be a nursery area due to the substrate type. Here larvae transported by the Caribbean current could settle and metamorphose into juveniles [27,35,66]. In their study on coral reef connectivity throughout the Caribbean, Schill et al. [58] stated that coral reefs rely heavily on ocean currents in order to maintain genetic vigour as currents are a valid source of new recruits which influences biomass, population persistence, resilience and species diversity. Sample location 3, based on its position, is directly impacted by westerly flowing currents and incidentally displayed the lowest average for total heterozygosity (0.736). This corroborates the indication that exploitation, genetic drift and potential larval retention may promote genetic differentiation among groups [67]. Additionally, Stoner et al. [62] cautioned that increased recruitment from upstream areas does not guarantee high queen conch production as mortality rates in newly settled invertebrates can be high. The highest juvenile densities are found on the southeastern edge of the bank [35,61], however the presence of small juveniles with limited capacity for migration and dispersal across Pedro Bank indicates that significant larval recruitment and settlement are occurring throughout the bank. It also suggests that replenishment of queen conch is not solely dependent upon recruitment from traditional nursery areas [11], that is, Pedro Bank may be self-recruiting [56] but still receives larvae from upstream areas which have greatest influence at the eastern end of the bank.

### 4.3 Management Implications

Over the years the impacts of commercial fishing and overexploitation of *L. gigas* on Pedro Bank have received considerable attention from a wide variety of stakeholders ranging from within Jamaica and the wider Caribbean, to international trading partners concerned with conservation of the species [3,7,68–70]. Solutions to these impacts have been sought through biological surveys and assessments [11,18,35,61] marine spatial planning [60] and genetic assessment [56].

However, management plans thus far have only utilized biological surveys to guide the queen conch fishery in Jamaica which is primarily based on Pedro Bank. The results from this study revealed a weak but significant population structure which may be habitat dependent and would benefit from EBM as suggested by Morris [18].

Genetic diversity of queen conch on Pedro Bank embodies that of a fit population in all monitoring zones sampled under prevailing environmental conditions, however the central to eastern and southeastern end of the bank comprising monitoring zones 1 and 3 are heavily fished and incidentally comprise dense patches of seagrass beds which potentially act as nursery areas for recruitment to the fishery. These zones are predominantly < 20 m in depth hence the increased exploitation as it is more economical and safer for commercial fishers to fill their quotas in these areas [35]. The deeper areas in open water locations are also more exposed to high waves [71] and ocean currents which are variable in strength [56]. The constant removal of species from one area can be detrimental as entire cohorts may be removed over successive fishing seasons causing the fishery to eventually collapse by the removal of reproductive individuals before they make a contribution to the fishery. This type of serial dilution was observed in the Bahamian queen conch fishery by Stoner et al. [71], to cause low densities of adult conch incapable of successful mating and reproduction hence a fishery on the verge of collapse if strong management measures are not instituted. Even though the total allowable catch for queen conch in Jamaica is relatively low per fishing season, commercial fishing should be encouraged across the entire Pedro Bank perhaps with incentives to offset economic losses such as an increased quota to fish in particular areas, thus safeguarding the genetic diversity across the bank. Instituting the marine spatial planning (MSP) for Pedro Bank as suggested by Baldwin et al. [60], would also allow for important nursery areas to be protected and permit the Fisheries Division to meet one objective of the conch management plan for rehabilitation of overexploited stocks.

A closed season to protect the reproductive period is a common management strategy in fisheries. This is particularly important for queen conch because they are susceptible to overexploitation when they aggregate in particular areas to spawn [2,52,72]. Reproduction in queen conch occurs year round in some areas [59,73], although reproduction seems to be triggered by temperature [18,21], so spawning tends to occur in the warmer months of the year [69,74]. The closed season for queen conch in Jamaica is set by the Fisheries Division through the Minister with responsibility of MICAF and currently varies from year to year. With the event of climate change and other environmental factors that may impact spawning, there needs to be a concerted effort to investigate temporal variability of the life history stages of *L. gigas* and as such institute appropriate periods for fishing restrictions suitable to the biology of the species. An alternate management measure to closed seasons may be closed areas as monitoring zones can be closed for specific periods to achieve the objectives of a closed season.

Without proper enforcement, management strategies are mere suggestions for good practices. Aiken et al. [13] has lamented the fact that enforcement has historically been a weak point for the fishing industry in Jamaica. Cochrane and Garcia [72] made the realistic observation that if enforcement is to take place in remote areas which cover a large expanse such as Pedro Bank the practicalities and cost of enforcement can be prohibitive. Although enforcement in Jamaica has been a challenge, exceptions to this are highlighted to some degree for the conch fishery because it is a commercial fishery and production is primarily export oriented. Whilst landings geared towards export are captured in the queen conch production for Jamaica, landings by artisanal fishermen for local consumption by restaurants and hotels may go unnoticed and hence not captured in the total estimated figure for queen conch production. In addition to contributing to serial dilution of stocks, this may affect the NTAC set by fisheries managers and hence calls for increased monitoring of harvesting areas and local landings by artisanal fishermen.

The identification of two possible subpopulations on Pedro Bank will assist fisheries managers in regulating harvesting areas and strengthen the certification scheme implemented to facilitate trade with the EU as entire sections of the bank may be closed off to fishing if necessary, and traceability of the individuals landed will be possible. The genetic information elucidated from the study will also strengthen the non-detrimental findings (NDF) required by CITES as traceability of the products can be substantiated along with the data provided for export. Knowing the genetic composition of the stocks on Pedro Bank and being able to trace the final product to the source stock will also serve as a deterrent to IUU fishing [75], which can only be substantially decreased with enforcement.

## 5. CONCLUSION

The results from this study revealed a weak but significant population structure for *L. gigas* on Pedro Bank as was seen in previous studies done elsewhere in the Caribbean [21,23,36,54, 67]. Queen conch displayed a patchy distribution across the monitoring zones sampled with significant but weak genetic variation found in zone 3 which is located on the heavily exploited eastern end of the bank. This genetic differentiation may be attributed to two main factors, namely the serial dilution of species and the complex oceanic current patterns associated with the eastern end of the bank. Pedro Bank is directly impacted by the westward flow of the Caribbean current and could serve as the primary recruitment area of queen conch larvae from upstream locations. The geographical and ecological conditions on the eastern end of the bank are also conducive to post larval settlement, however the high densities of juvenile conch found throughout the bank coupled with high gene flow across the bank suggests that self-recruitment is taking place and is possible due to the different depth strata presently unexploited within the monitoring zones. These recruitment options available to the *L. gigas* stocks on Pedro Bank should ensure a viable population continues to exist with the use of proper management tools incorporating all available data into ecosystem management and continued monitoring of the stocks.

## 6. ACKNOWLEDGMENT

I wish to thank the Fisheries Division currently the National Fisheries Authority in the Ministry of Industry, Commerce and Agriculture, for their continued support throughout the project, especially Mr. Stephen Smikle. I am also grateful to the “One Leaders Diving Team”, in particular, Mr. Junior Squire and Mrs. Kimberly Cooke-Panton for their assistance in sample collection. Special thanks to my supervisors who provided academic stimulation and advice on numerous occasions. My sincere gratitude to the Link Foundation through the Smithsonian Institution which provided the opportunity to conduct research at the Smithsonian Marine Station (SMS).

## 7. AUTHORS CONTRIBUTION

Conceptualization: AB, MKW, KAA

Formal Analysis: AB, NT, IS

Funding Acquisition: KAA, SB, MKW

Investigation: AB

Methodology: NT, IS, AB

Project Administration: KAA, MKW, AB

Resources: AB, IS, NT

Supervision: KAA, MKW, MER

Validation: IS, NT, AB

Writing – Original Draft: AB

Writing – Review & Editing: AB, KAA, MKW, IS, MER, SB

